# Assessing ambitious nature conservation strategies within a 2 degree warmer and food-secure world

**DOI:** 10.1101/2020.08.04.236489

**Authors:** Marcel T.J. Kok, Johan R. Meijer, Willem-Jan van Zeist, Jelle P. Hilbers, Marco Immovilli, Jan H. Janse, Elke Stehfest, Michel Bakkenes, Andrzej Tabeau, Aafke M. Schipper, Rob Alkemade

## Abstract

Global biodiversity is projected to further decline under a wide range of future socio-economic development pathways, even in sustainability oriented scenarios. This raises the question how biodiversity can be put on a path to recovery, the core challenge for the CBD post-2020 global biodiversity framework. We designed two contrasting, ambitious global conservation strategies, ‘Half Earth’ (HE) and ‘Sharing the Planet’ (SP), and evaluated their ability to restore terrestrial and freshwater biodiversity and to provide ecosystem services while also mitigating climate change and ensuring food security. We applied the integrated assessment framework IMAGE with the GLOBIO biodiversity model, using the ‘Middle of the Road’ Shared Socio-economic Pathway (SSP2) with its projected human population growth as baseline.

We found that both conservation strategies result in a reduction in the loss of biodiversity and ecosystem services globally, but without additional measures to achieve effective climate mitigation they will be insufficient to restore biodiversity. The HE strategy performs better for terrestrial biodiversity protection (biodiversity intactness (MSA), Red List Index, geometric mean abundance) in currently still natural regions, reflecting global conservation priorities. The SP strategy yields more improvements for biodiversity in human-used areas, aquatic biodiversity and for regulating ecosystem services (pest control, pollination, erosion control), reflecting regional priorities. However, ‘conservation only’ scenarios show a considerable increase in food security risks (especially in Sub-Saharan Africa) compared to the baseline and limited reduction of global temperature increase. Only when conservation strategies are combined with climate change mitigation efforts and additional actions especially in the agricultural and energy system into a portfolio of ‘integrated sustainability measures’, both conservation strategies result in restoring biodiversity to current values or even some improvement, while keeping global warming below two degrees and keeping food security risks below baseline. Minimizing food wastes and reducing consumption of animal products will be crucial.

## 1. Introduction

The decline of biodiversity continues despite global commitments under the UN Convention on Biological Diversity (CBD). In 2020 the CBD Strategic Plan 2010-2020 will come to an end and countries have to renew their commitments. In view of limited progress in realising CBD goals and targets (Tittensor et al., 2014; IPBES, 2019, sCBD, 2020), countries need to design and agree upon a new and more effective global biodiversity framework for the coming decades to realise the 2050 vision of the CBD. This new framework needs to be part and parcel of the broader sustainable development agenda, especially the internationally agreed Agenda 2030 and its Sustainable Development Goals (SDGs) (UN GA, 2015). Agenda 2030 emphasizes the need to develop biodiversity conservation strategies that are consistent with other SDGs – and sustainable development in general – reflecting the increasing acknowledgement of the importance of biodiversity and natures contributions to people for human well-being (Blicharska et al., 2019; IPBES, 2019; Lucas et al., 2014).

A variety of scenario-based analyses and modelling approaches (IPBES, 2016) can help to understand future impacts of drivers of biodiversity loss and to explore the potential of various measures to achieve global goals and targets. Current explorative scenario-projections, covering a wide range of plausible socio-economic pathways, indicate that global biodiversity will continue to decline, even under more optimistic scenarios oriented towards sustainability (Shin et al., 2019; Schipper et al. 2020; Pereira et al., 2020). Moreover, despite the 2015 Paris Agreement to limit global warming to well-below 2 degrees, the world is currently heading towards a global temperature increase of 3.2 degrees (UNEP, 2019), and more than 800 million people are still food insecure (FAO, 2019). This raises the question of how to put nature on a path to recovery, while also halting climate change and contribute to ending hunger and feeding a growing and wealthier global population.

To answer this question, scenario analysis recently shifted to consider ambitious objectives for nature conservation and desirable nature futures (Rosa et al., 2017; Pereira et al., 2020). Recent papers have assessed what efforts are necessary to put nature on a path to recovery (Mace et al., 2018; Leclere et al., 2018, 2020 (accepted); Tickner et al., 2020), building on earlier work to halt the loss of biodiversity and meet the Aichi targets (van Vuuren et al., 2015; Kok et al., 2018; Visconti et al., 2016). Challenges remain, however, to include all relevant drivers of biodiversity loss, apply multiple perspectives to area-based conservation and to integrate biodiversity outcomes with other relevant SDGs, while considering multiple visions for nature (Rosa et al., 2020). Furthermore, efforts need to be put to examine how nature supports socio-economic development and human well-being (Rosa et al., 2017). Ultimately, there is a need to understand which ambitious long-term conservation strategies can restore nature and contribute to meeting other societal goals, such as, in this analysis, food security and climate objectives (Allan et al., under review; IPBES, 2019, p. 56; Rosa et al., 2017).

We aim to fill these gaps by applying a solution-oriented, model-based scenario-analysis to evaluate two alternative, ambitious conservation strategies, labelled ‘Half Earth’ and ‘Sharing the Planet’. We evaluate both strategies for their realisation of internationally agreed goals for biodiversity (the 2050 Vision and the emerging post-2020 global biodiversity framework being developed under the CBD), climate change (UNFCCC Paris Agreement) and food security (SDG 2 End hunger).

In response to recent calls, our scenarios include multiple perspectives on conservation (Bhola et al., 2020; Immovilli and Kok, 2020) and values of nature (Pascual et al., 2017; Pereira et al., 2020; Rosa et al., 2017; van Zeijts et al., 2017). Including these different perspectives helps to show how different valuations of nature may be reflected in different conservation strategies and pathways. This in turn, also helps to show how different approaches to conservation can become contested not only between conservation points of view, but also from for example food security and equity perspectives (Ellis and Mehrabi, 2019; Mehrabi et al., 2018, Otero et al., 2020; Schleicher et al., 2019).

The ‘Half Earth’ (HE) strategy emerges from a focus on protecting nature for its intrinsic value. Several proposals have been made for ambitious area-based conservation efforts, including conserving half of the earth’s surface (Wilson, 2016; Pimm et al., 2018), nature needs half (Locke, 2013) and the global deal for nature (Dinerstein et al., 2017, 2019). Despite differences in methods applied, proponents agree on the need to promote wilderness and separating human influence from nature as the most important solution to halt biodiversity loss. To achieve such bold targets, these approaches rely on extending protected areas, conserving other remaining natural areas and restoration efforts as their main strategies, while recognizing that the strategy would be applied differentially to different regions (Locke et al., 2019). A consequence of setting aside large areas for nature conservation is that it limits space for producing food and other agricultural and forestry products, requiring massive improvement of agricultural productivity.

The ‘Sharing the Planet’ strategy is underpinned by the principle of ‘living with nature’ (Hinchliffe, 2006; Turnhout et al., 2013) and entails both instrumental and relational valuations of nature. Conservation is ‘convivial’ insofar as it aims to maintain biodiversity and create value at a local level, it does not seek to separate humans from nature and it addresses structural social inequalities and injustices (Büscher & Fletcher, 2020). Protecting and supporting nature’s contributions to people (NCP) are therefore the main objectives of this perspective. With this regard, proposals have been made to prioritize the multifunctionality of a landscape and the delivery of multiple benefits through ecosystem services. Thus, biodiversity conservation is one of the many benefits a landscape can provide, along with others such as food security, poverty alleviation, sense of identity, etc. (Kremen & Merenlender, 2018; Perfecto & Vandermeer, 2016). Agriculture production is part of the integrated and productive landscapes.

The impacts of both HE and SP conservation strategies on biodiversity and food security are evaluated in scenarios where climate change mitigation is consistent with currently committed global mitigation efforts (expected to result in a global warming of 3.2 °C in 2100 (UNEP, 2019) and in scenarios where global warming is limited to 2 °C and thus additional sustainability measures in the food and agricultural system are applied. To the former we refer as ‘conservation only’ scenarios, to the latter as ‘integrated sustainability’ scenarios. The scenarios are evaluated against, the ‘Middle of the Road’ scenario of the Shared Socio-economic Pathways (SSP2) (Riahi et al., 2017). SSP2 describes a world in which social, economic, and technological trends do not shift markedly from historical patterns. We project the scenarios to 2070 and report results also for two intermediate years (2030 and 2050). We do not address the governance needed to realise these strategies, nor does the model-framework allow us to explore feedbacks of these strategies on economy, production systems or population (see Rosa et al., 2017, 2020). The integrated assessment model IMAGE (Stehfest et al., 2014) and the global biodiversity model GLOBIO (Schipper et al., 2020; Janse et al., 2015) are applied to evaluate the outcomes of these pathways in terms of various metrics of biodiversity, ecosystem services, climate change and food security and to assess the efforts needed for this.

## 2. Methods

### 2.1 Conservation strategies

As a starting point of the scenario analysis, we elaborated consistent narratives for two contrasting global conservation strategies, ‘Half Earth’ and ‘Sharing the Planet’, based on literature review and recent debates in the context of CBD’s post-2020 global biodiversity framework (see S.1 and Immovilli and Kok, 2020 for more elaborate descriptions, see also Bhola et al., 2020).

In the ‘Half Earth’ strategy, nature is valued for its intrinsic value. The concept of wilderness and naturalness are central to this strategy, to be protected from human pressures (Kopnina, 2016; Wuerthner, 2014). In order to retain the naturalness of ecosystems, Protected Areas (PAs) and other area-based conservation measures are cornerstones of this strategy. Ecosystem services and natures contributions to people are considered as a co-benefit, but not prioritized. This conservation strategy requires a land sparing approach for agriculture, based on sustainable intensification aiming at closing the yield gaps (Garnett et al., 2013; Phalan et al., 2016). This type of intensification draws upon wide technological developments and innovations, such as more efficient irrigation and nutrient use, pest management and genetic modification, and aims for eco-efficiency and reduction of externalities. There is less reliance on bio-energy. Also other pressures besides agriculture (like hydropower) are restricted in the protected half. In river basins, new hydropower dams would be only allowed in the ‘non-protected half’, while in the ‘protected half’ measures will be taken to partly restore the flow of water and reduce river fragmentation.

In the ‘Sharing the Planet’ strategy, conservation measures that support and enhance the provision of ecosystem services and nature’s contributions to people are prioritized. This strategy premises on the concept of ‘living with nature’ (Turnhout et al., 2013; Buscher and Fletcher, 2020). In agriculture, the dominant landscape is a heterogenous mosaic, comprising a combination of natural habitat patches and agriculture, a so-called agro-ecological matrix, resulting in improved provision of ecosystem services (Kremen & Merenlender, 2018; Perfecto & Vandermeer, 2016). Food production, carbon storage, pollination, water and nutrient retention, biodiversity conservation and other services are achieved within the mosaic landscape following the path of ecological intensification. This type of intensification is concerned with the optimization of ecosystem benefits and it primarily relies on ecological knowledge, labor-intensive or smart mechanization systems, as applied in agroecology, organic farming, agroforestry and diversified farming systems (Tittonel, 2014). There is less reliance on bio-energy, and additional hydropower capacity is only possible while respecting ecological flow requirements and minimizing further river fragmentation.

### 2.2 Defining scenarios

The Half Earth (HE) and Sharing the Planet (SP) strategies were translated into four quantitative scenarios (Table 1), including ‘conservation only’ scenarios (HE-co and SP-co) and ‘integrated sustainability’ scenarios combining conservation with measures to limit global warming to 2 °C and additional sustainability measures in the food and agricultural system (HE-is and SP-is). We created global maps of potential conservation areas consistent with the HE and SP narratives as the starting point of the analysis (see S. 2 for further details).

**TABLE 1.**
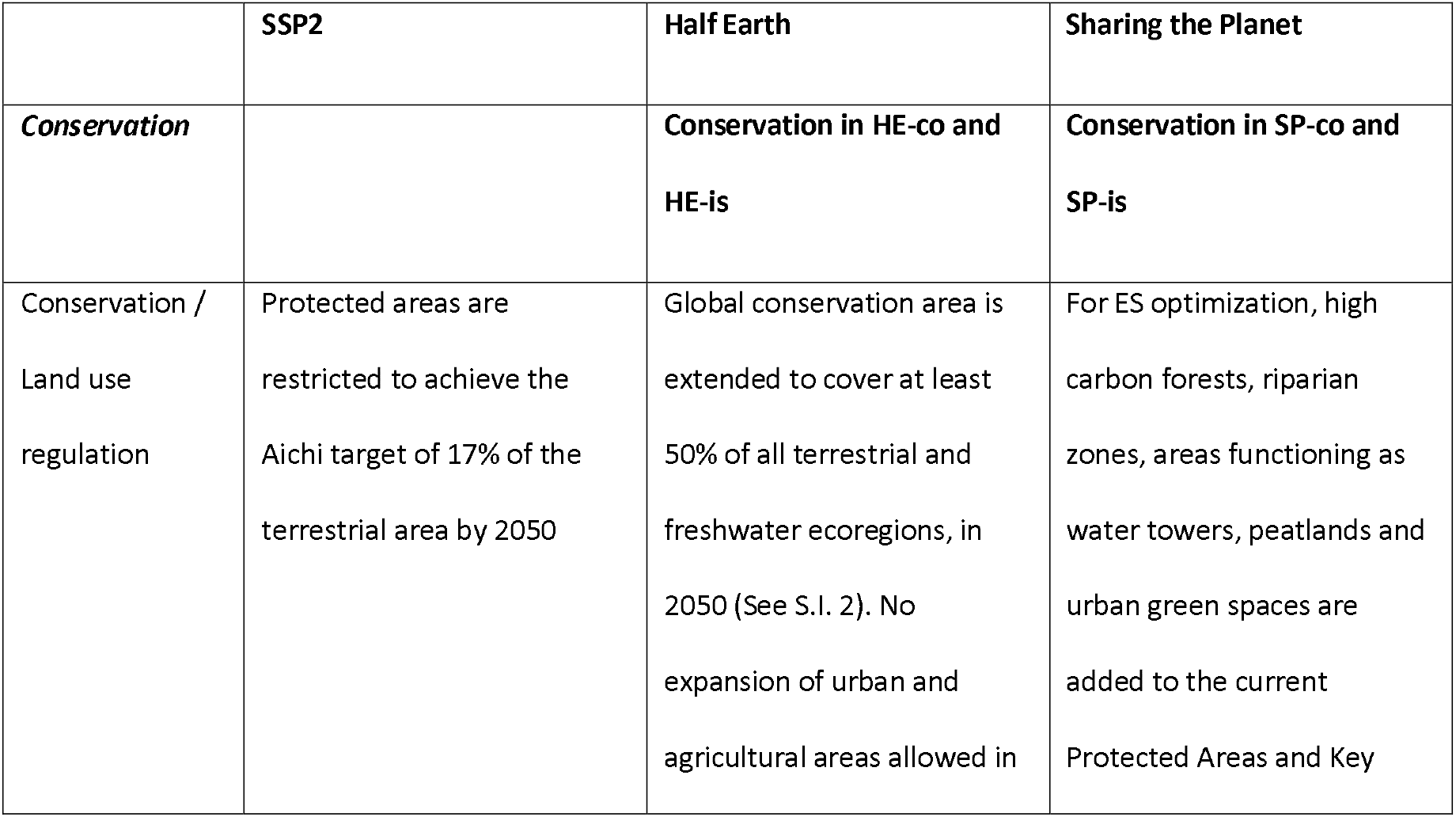
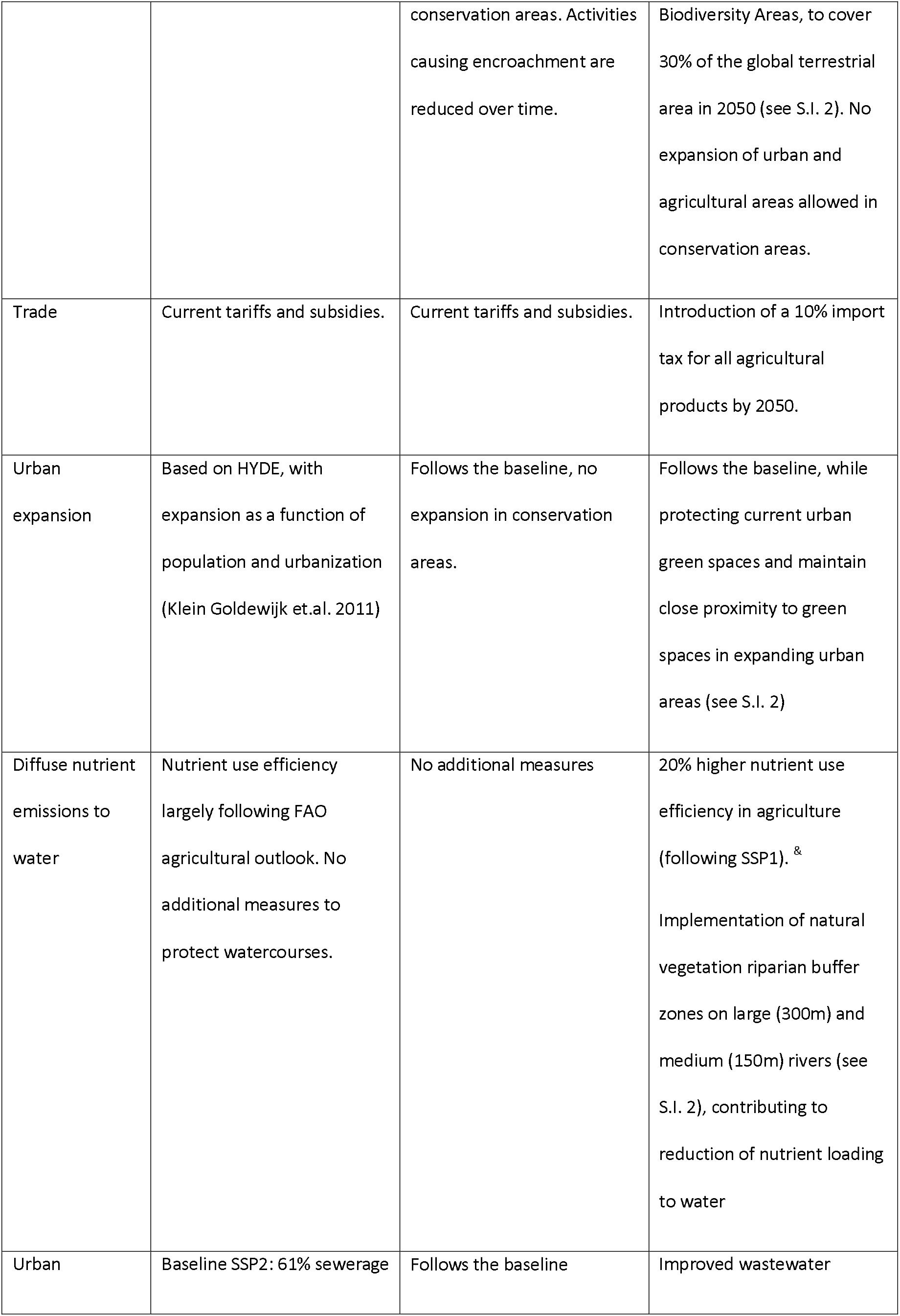
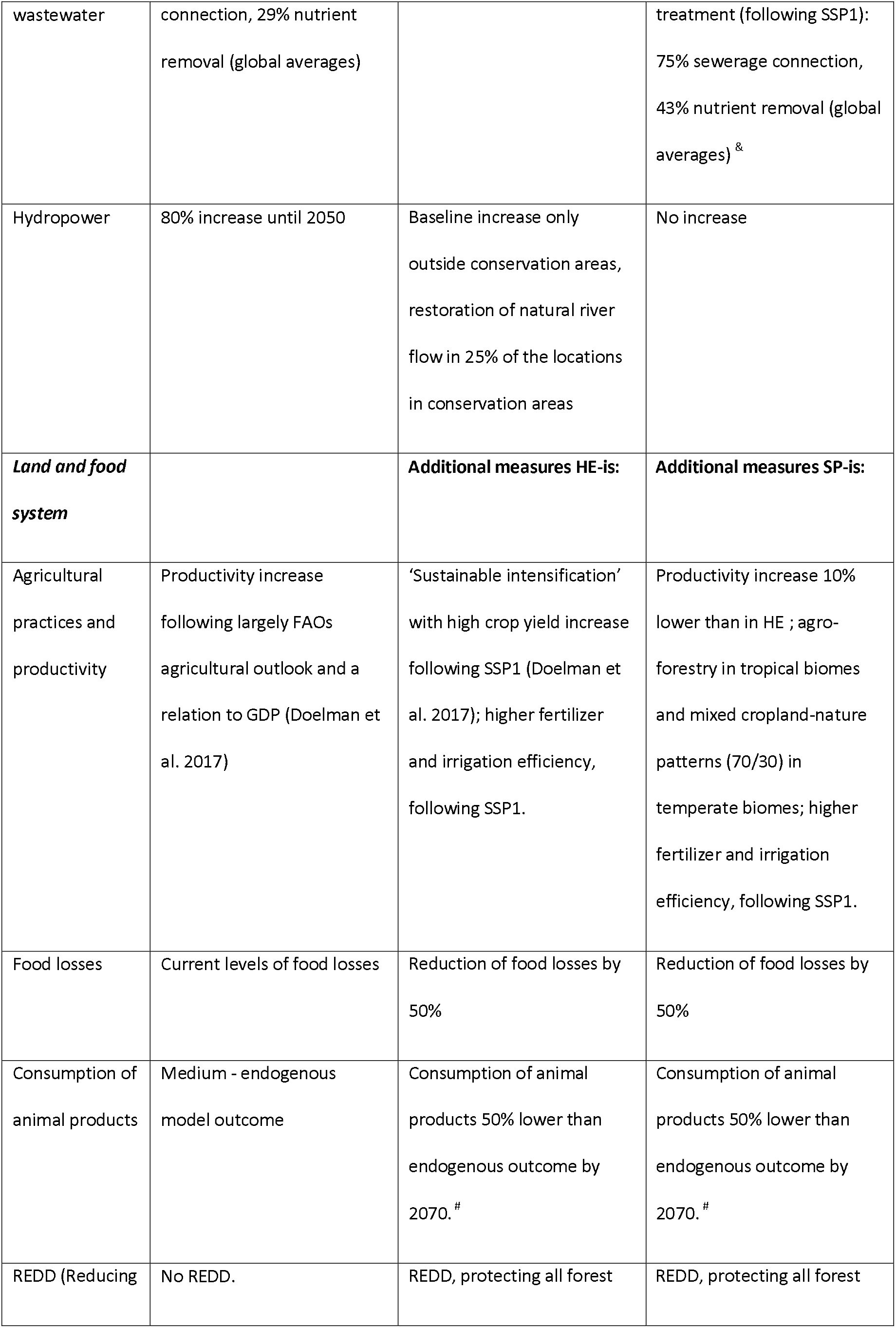
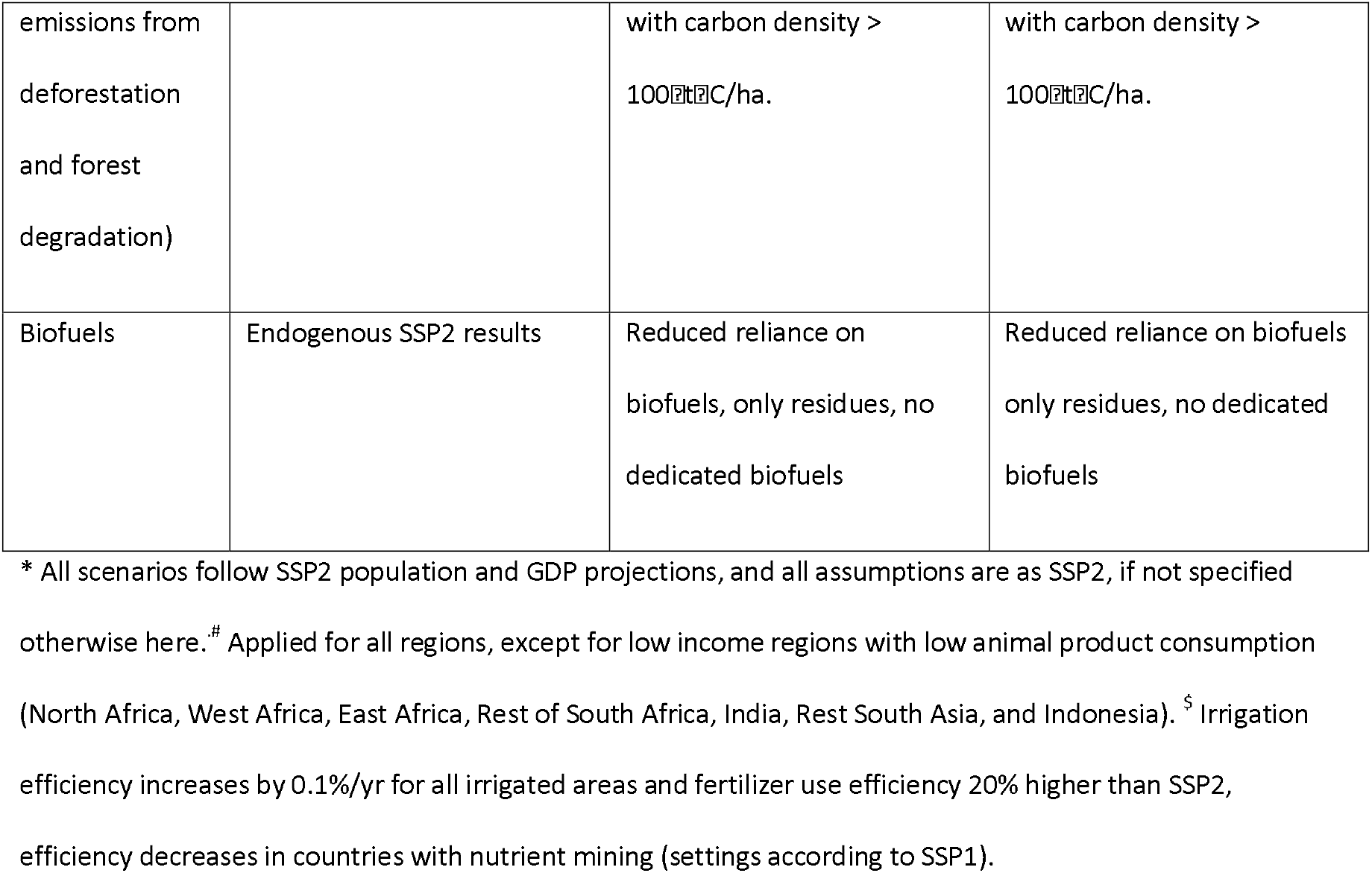
Key assumptions in the SSP2 baseline and the four scenarios*

|  | SSP2 | Half Earth | Sharing the Planet |
| --- | --- | --- | --- |
| *Conservation* |  | Conservation in HE-co and HE-is | Conservation in SP-co and SP-is |
| Conservation /<br>Land use<br>regulation | Protected areas are restricted to achieve the Aichi target of 17% of the terrestrial area by 2050 | Global conservation area is extended to cover at least 50% of all terrestrial and freshwater ecoregions, in 2050 (See S.I. 2). No expansion of urban and agricultural areas allowed in | For ES optimization, high carbon forests, riparian zones, areas functioning as water towers, peatlands and urban green spaces are added to the current Protected Areas and Key |
|  |  | conservation areas. Activities causing encroachment are reduced over time. | Biodiversity Areas, to cover 30% of the global terrestrial area in 2050 (see S.l. 2). No expansion of urban and agricultural areas allowed in conservation areas. |
| Trade | Current tariffs and subsidies. | Current tariffs and subsidies. | Introduction of a 10% import tax for all agricultural products by 2050. |
| Urban expansion | Based on HYDE, with expansion as a function of population and urbanization (Klein Goldewijk et.al. 2011) | Follows the baseline, no expansion in conservation areas. | Follows the baseline, while protecting current urban green spaces and maintain close proximity to green spaces in expanding urban areas (see S.l. 2) |
| Diffuse nutrient emissions to water | Nutrient use efficiency largely following FAO agricultural outlook. No additional measures to protect watercourses. | No additional measures | 20% higher nutrient use efficiency in agriculture (following SSP1).<sup>&</sup><br>Implementation of natural vegetation riparian buffer zones on large (300m) and medium (150m) rivers (see S.l. 2), contributing to reduction of nutrient loading to water |
| Urban | Baseline SSP2: 61% sewerage | Follows the baseline | Improved wastewater |
| wastewater | connection, 29% nutrient removal (global averages) |  | treatment (following SSP1):<br>75% sewerage connection,<br>43% nutrient removal (global averages) & |
| Hydropower | 80% increase until 2050 | Baseline increase only<br>outside conservation areas,<br>restoration of natural river<br>flow in 25% of the locations<br>in conservation areas | No increase |
| ***Land and food system*** |  | **Additional measures HE-is:** | **Additional measures SP-is:** |
| Agricultural practices and productivity | Productivity increase following largely FAOs agricultural outlook and a relation to GDP (Doelman et al. 2017) | 'Sustainable intensification' with high crop yield increase following SSP1 (Doelman et al. 2017); higher fertilizer and irrigation efficiency, following SSP1. | Productivity increase 10% lower than in HE ; agro-forestry in tropical biomes and mixed cropland-nature patterns (70/30) in temperate biomes; higher fertilizer and irrigation efficiency, following SSP1. |
| Food losses | Current levels of food losses | Reduction of food losses by 50% | Reduction of food losses by 50% |
| Consumption of animal products | Medium - endogenous model outcome | Consumption of animal products 50% lower than endogenous outcome by 2070. # | Consumption of animal products 50% lower than endogenous outcome by 2070. # |
| REDD (Reducing | No REDD. | REDD, protecting all forest | REDD, protecting all forest |
| emissions from<br>deforestation<br>and forest<br>degradation) |  | with carbon density ><br>100 tC/ha. | with carbon density ><br>100 tC/ha. |
| Biofuels | Endogenous SSP2 results | Reduced reliance on<br>biofuels, only residues, no<br>dedicated biofuels | Reduced reliance on biofuels<br>only residues, no dedicated<br>biofuels |

\* All scenarios follow SSP2 population and GDP projections, and all assumptions are as SSP2, if not specified otherwise here.<sup>#</sup> Applied for all regions, except for low income regions with low animal product consumption (North Africa, West Africa, East Africa, Rest of South Africa, India, Rest South Asia, and Indonesia).<sup>§</sup> Irrigation efficiency increases by 0.1%/yr for all irrigated areas and fertilizer use efficiency 20% higher than SSP2, efficiency decreases in countries with nutrient mining (settings according to SSP1).

To assess the impacts of ‘conservation only’ strategies on biodiversity and food security, we evaluated the HE and SP strategies in a situation where climate change is hardly mitigated (RCP 6.0) and in which no further measures in the food and agricultural system are taken. The RCP 6.0 scenario is close to the SSP2 baseline result at a radiative forcing of 6.0 W m-2, and is consistent with currently committed global mitigation efforts (UNEP, 2019). Next, we combined the HE and SP conservation strategies with measures to limit global warming to 21°C (RCP 2.6; Van Vuuren et al., 2011) and measures in the food and agricultural system needed to safeguard global food security. These ‘integrated sustainability’ pathways combine area-based conservation and restoration with changes in land use, agricultural production and food consumption to mitigate climate change (e.g., reducing food waste, reducing animal product consumption and further increasing agricultural productivity). We adjusted some assumptions regarding the energy system compared to the standard SSP2, RCP 2.6 energy-mix, notably with respect to biomass and hydropower, to take into account biodiversity concerns (Table 1).

### 2.3 IMAGE and GLOBIO models and indicators

We applied the integrated assessment model IMAGE (Stehfest et al., 2014) and the biodiversity model GLOBIO (Schipper et al., 2020, Janse et al., 2015) to evaluate the four scenarios. IMAGE assesses global environmental change under future socio-economic development scenarios. It includes a set of connected models describing the energy, the agricultural economy and land use, natural vegetation and the climate system. For our analysis we particularly used the MAGNET model for calculating changes in agricultural demand, production and trade (Woltjer et al., 2014); relied on earlier calculations of TIMER for changes in the energy system and the demand for bioenergy and hydropower (van Vuuren, 2007); the GNM model for calculating nutrient loads in water systems (Beusen et al., 2015) and LPJmL for carbon, water, crop and grass yields, and vegetation dynamics (Schaphoff et al., 2018, Müller et al. 2016); the GISMO model to asses impacts on food security (Lucas et al., 2019). Socio-economic processes, especially in MAGNET and TIMER, distinguish 26 world regions, while land use and all biophysical processes are resolved at a 5 x 5 arcminutes and 30 x 30 arcminutes resolution.

The GLOBIO model includes four main components: 1) GLOBIO for assessing terrestrial biodiversity intactness (Schipper et al., 2020), 2) GLOBIO-Aquatic for assessing freshwater biodiversity intactness (Janse et al., 2015), 3) GLOBIO-Species for assessing the distribution and abundance of vertebrate species, based on the INSIGHTS modelling framework (Visconti, 2016; Baseiro, 2020, see also S. 3), and 4) GLOBIO-ES for assessing ecosystem services (Schulp et al., 2012). Indicators assessed with GLOBIO are a function of multiple human pressures: land use, fragmentation, road disturbance, encroachment by hunting, climate change, atmospheric nitrogen deposition, nutrient emissions to water and hydrological disturbance. IMAGE scenario outcomes are used to quantify various of these pressures. We downscaled changes in land use calculated in the IMAGE model downscaled to a 10 arcseconds resolution (~300×300meter at the equator) by the land use allocation module integrated in GLOBIO (Schipper et al., 2020). GLOBIO also uses estimates on global temperature change, fertilizer use and nitrogen deposition from IMAGE. Nutrient loads to water were calculated by GNM (Beusen et al., 2015) and river discharge by LPJmL (Biemans et al., 2011) and PCR-GLOBWB (Sutanadjaja et al., 2018). Road infrastructure data were used from the GRIP database (Meijer et al., 2018).

Based on the indicators available in GLOBIO and IMAGE and following the proposal by Mace et al. (2018), we quantified three complementary biodiversity indicators representing trends in populations, extinctions and integrity (functional diversity) (Table 2): 1) the geometric mean abundance (GMA; Visconti et al. 2016), 2) the Red List Index (RLI; Butchart et al., 2007) and 3) the mean species abundance (MSA; Schipper et al. 2020; Janse et al. 2015). To cover ecosystem services or nature’s benefits to people, we include six terrestrial ecosystem services (crop production, grass & fodder production, pest control, pollination, carbon sequestration and erosion control) and four aquatic ones (water scarcity reduction, flood risk reduction, natural water purification and health and recreational value of lakes).

**TABLE 2.**
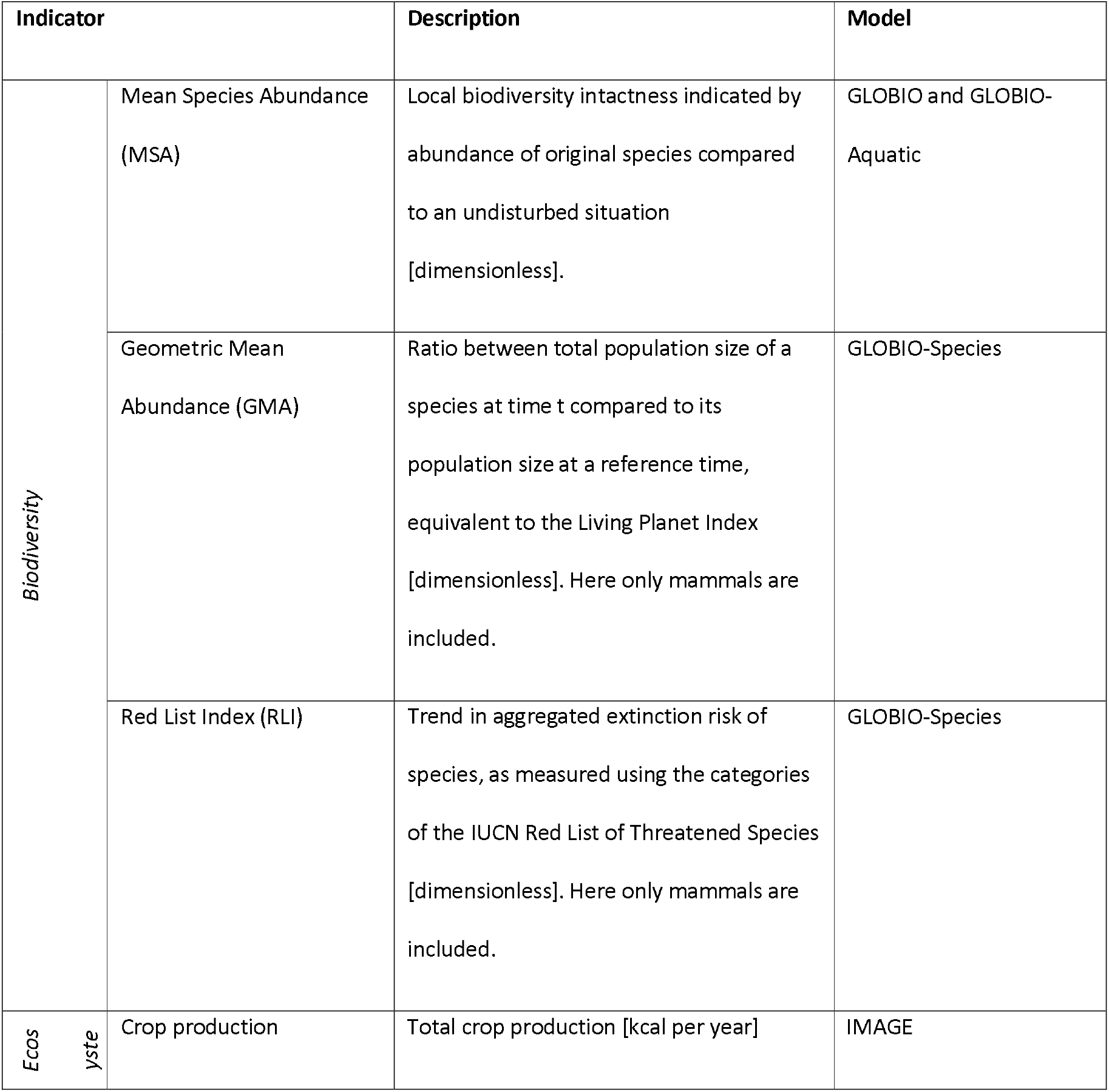
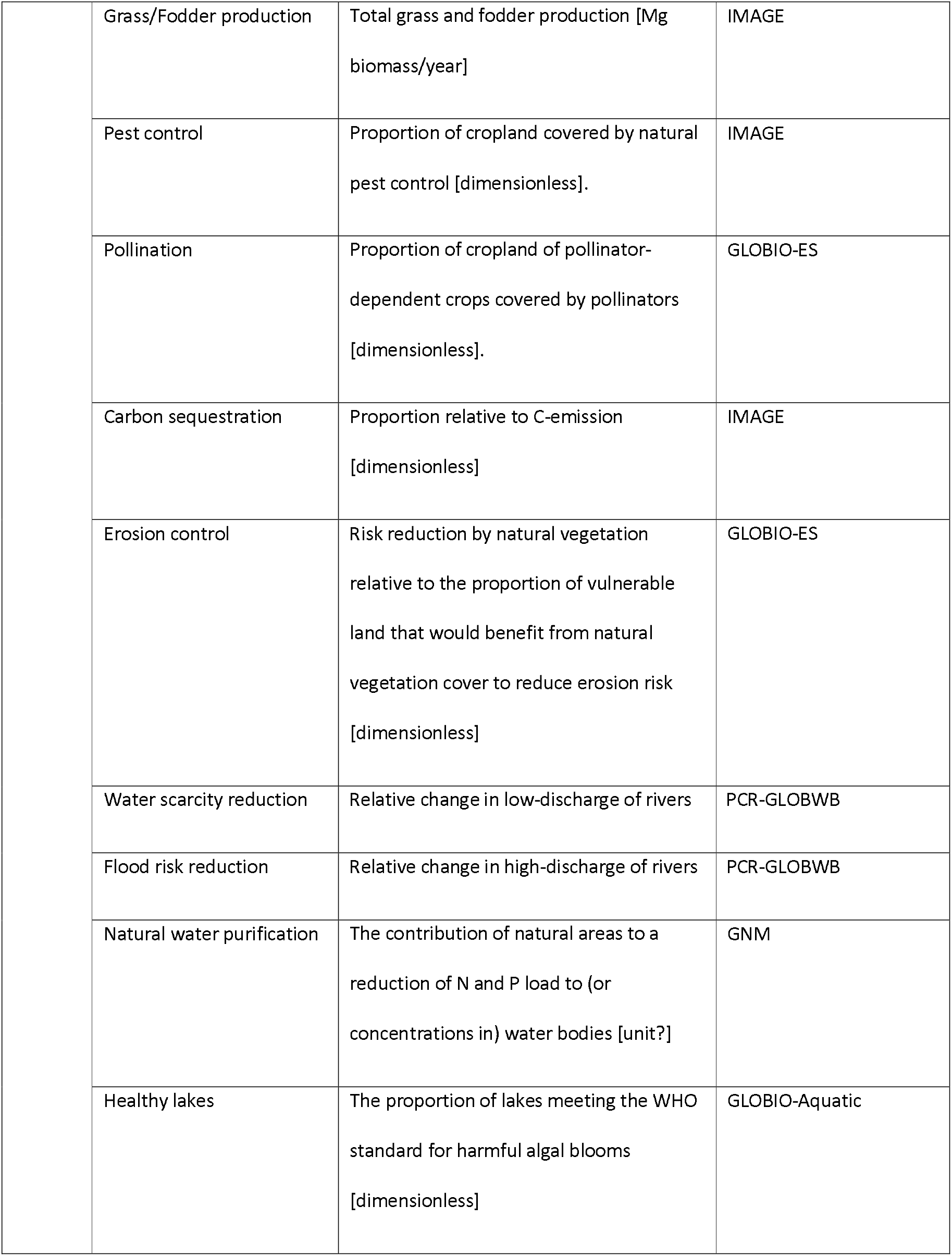
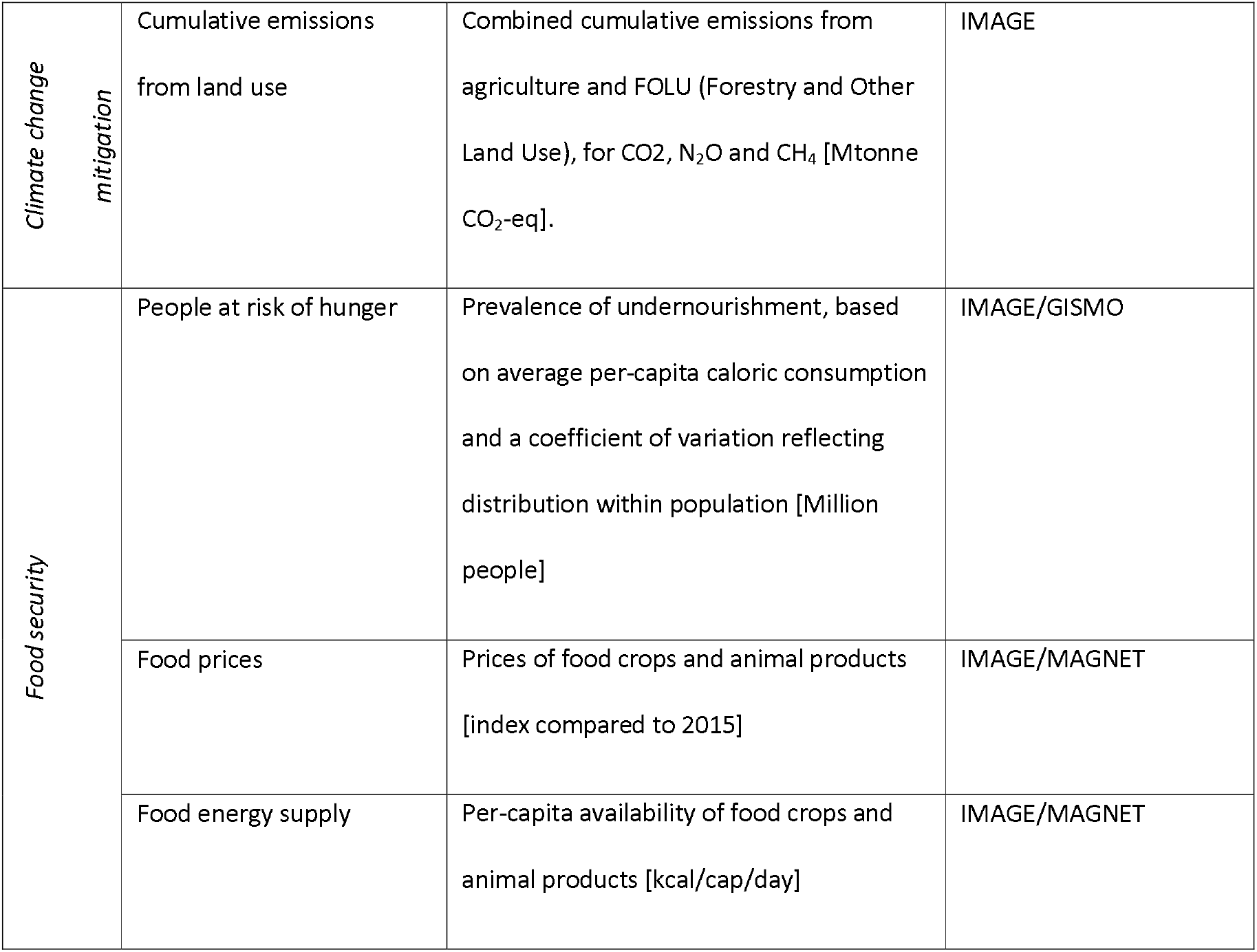
Overview of indicators (see also S. 4)

| Indicator |  | Description | Model |
| --- | --- | --- | --- |
| Biodiversity | Mean Species Abundance (MSA) | Local biodiversity intactness indicated by abundance of original species compared to an undisturbed situation [dimensionless]. | GLOBIO and GLOBIO-Aquatic |
|  | Geometric Mean Abundance (GMA) | Ratio between total population size of a species at time t compared to its population size at a reference time, equivalent to the Living Planet Index [dimensionless]. Here only mammals are included. | GLOBIO-Species |
|  | Red List Index (RLI) | Trend in aggregated extinction risk of species, as measured using the categories of the IUCN Red List of Threatened Species [dimensionless]. Here only mammals are included. | GLOBIO-Species |
| Ecosystems | Crop production | Total crop production [kcal per year] | IMAGE |

|  |  |  |
| --- | --- | --- |
| Grass/Fodder production | Total grass and fodder production [Mg biomass/year] | IMAGE |
| Pest control | Proportion of cropland covered by natural pest control [dimensionless]. | IMAGE |
| Pollination | Proportion of cropland of pollinator-dependent crops covered by pollinators [dimensionless]. | GLOBIO-ES |
| Carbon sequestration | Proportion relative to C-emission [dimensionless] | IMAGE |
| Erosion control | Risk reduction by natural vegetation relative to the proportion of vulnerable land that would benefit from natural vegetation cover to reduce erosion risk [dimensionless] | GLOBIO-ES |
| Water scarcity reduction | Relative change in low-discharge of rivers | PCR-GLOBWB |
| Flood risk reduction | Relative change in high-discharge of rivers | PCR-GLOBWB |
| Natural water purification | The contribution of natural areas to a reduction of N and P load to (or concentrations in) water bodies [unit?] | GNM |
| Healthy lakes | The proportion of lakes meeting the WHO standard for harmful algal blooms [dimensionless] | GLOBIO-Aquatic |
| *Climate change mitigation* | Cumulative emissions from land use | Combined cumulative emissions from agriculture and FOLU (Forestry and Other Land Use), for CO<sub>2</sub>, N<sub>2</sub>O and CH<sub>4</sub> [Mtonne CO<sub>2</sub>-eq]. | IMAGE |
| *Food security* | People at risk of hunger | Prevalence of undernourishment, based on average per-capita caloric consumption and a coefficient of variation reflecting distribution within population [Million people] | IMAGE/GISMO |
|  | Food prices | Prices of food crops and animal products [index compared to 2015] | IMAGE/MAGNET |
|  | Food energy supply | Per-capita availability of food crops and animal products [kcal/cap/day] | IMAGE/MAGNET |

As indicator for the impact on climate change we use the cumulative emissions from agriculture, forestry and other land use. Food security impacts were analysed in terms of food availability, food prices and people at risk of hunger. While food security has many more aspects, most global modelling studies rely on these indicators (Hasegawa et al. 2018, Van Meijl et al. 2020).

## 3. Results

### 3.1 Overall results for biodiversity and ecosystem services, climate change and food security

In this section we show the overall performance of the HE and SP pathways, compared to the baseline (Figure 1), while detailed results are presented in sections 3.2 through 3.4 and additional regional results in S.5.

In the baseline, environmental changes continue according to the current trends: land conversion will still be significant as shown by the increase of agricultural land and continued climate change. Due to scarcity of land and increased demand, global food prices will increase but the fraction of people at risk of hunger will decline. A continuous decline of biodiversity is expected and most regulating ecosystem services will decrease.

**FIGURE 1:**
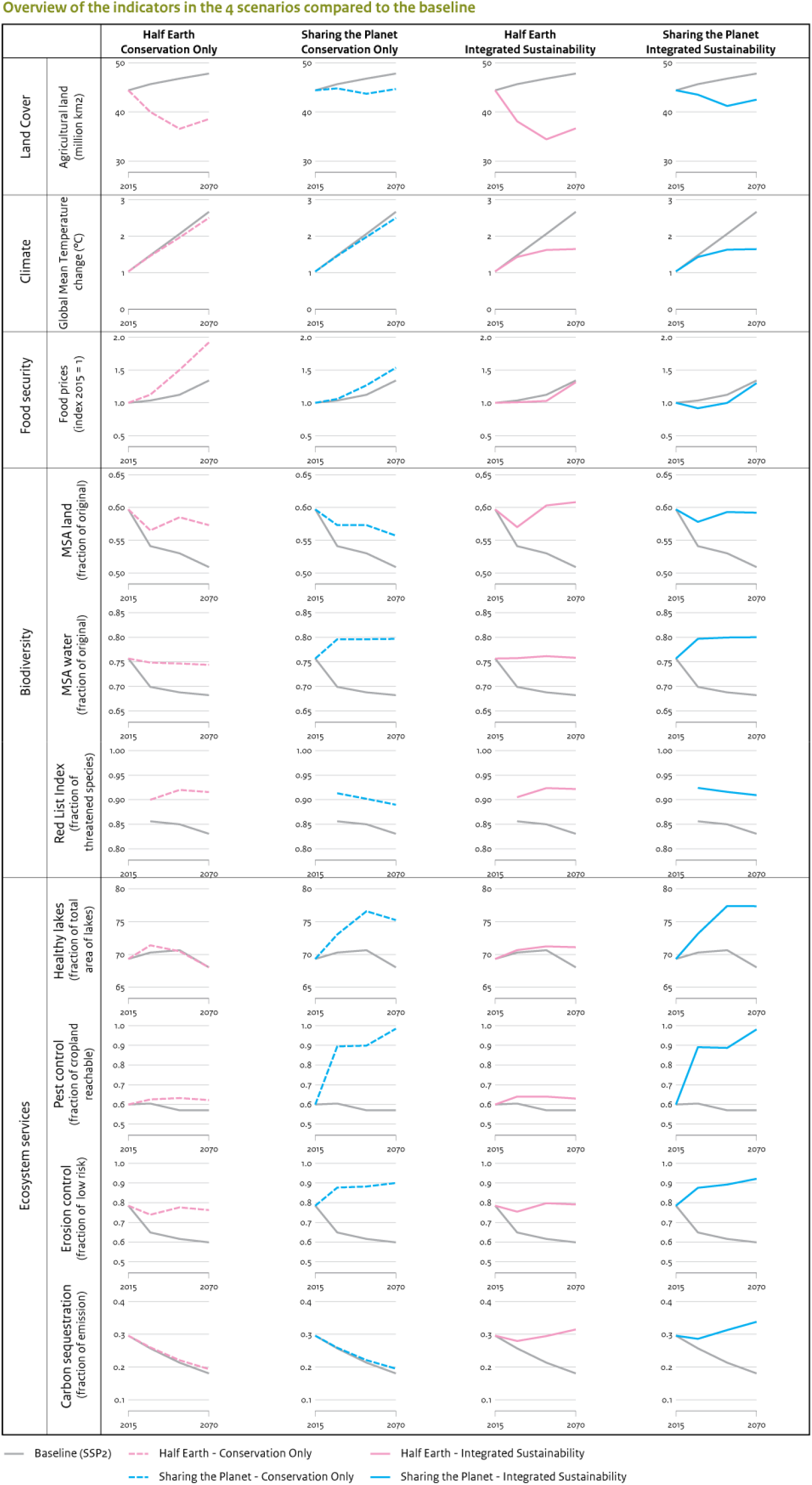
Overall global results for land use, climate change, food security, biodiversity and ecosystem services for selected indicators (Baseline, HE-co, HE-is, SP-co, SP-is; 2015-2070).

In the HE pathway, protecting 50% of the land surface combined with sustainable agricultural intensification results in substantial reduction of agricultural area. Implemented as ‘conservation only’ (HE-co), large benefits for biodiversity are expected and slightly better performance of regulating ecosystem services, but only limited climate benefits and an increase in food security risks.

The SP pathway, with lower levels of protection and agriculture moving towards agro-ecological approaches, shows less reduction in agricultural area and somewhat lower benefits for terrestrial but better results for aquatic biodiversity than the HE-pathway. Implemented as ‘conservation only’ (SP-co), limited climate benefits are expected and risks for food security still increase but less than in HE-co. SP performs substantially better for most ecosystem services as well as for aquatic biodiversity.

Introducing integrated sustainability measures, in both pathways (HE-is, SP-is), reduces food security risks and enables climate mitigation using land- and nature-based mitigation potential and creates further benefits for biodiversity and ecosystem services.

### 3.2 Land use change, climate change mitigation and food security

The baseline scenario (SSP2) shows a continued expansion of agricultural areas by 8% up to 2070 (Figure 2). In contrast, the HE scenarios, with substantial expansion of protected areas, show declines in cropland (5% and 17% for HE-co and HE-is respectively) and especially pasture (23% and 25% for HE-co and HE-is respectively) by 2050 and again an expansion after that. The SP scenarios show modest changes of agricultural land up to 2070 with mainly decrease of pastures in 2030-2050 (constant for SP-co and 4% decrease for SP-is), resulting from the intermediate position between the baseline and the HE pathway with respect to agricultural productivity and expansion of protected areas. The integrated sustainability measures in HE-is and SP-is stimulate agricultural productivity even more than in the baseline and reduce food demand, due to waste reduction and dietary changes. The recovery of mainly non-forest natural areas continues in both HE and SP until 2050.

**FIGURE 2:**
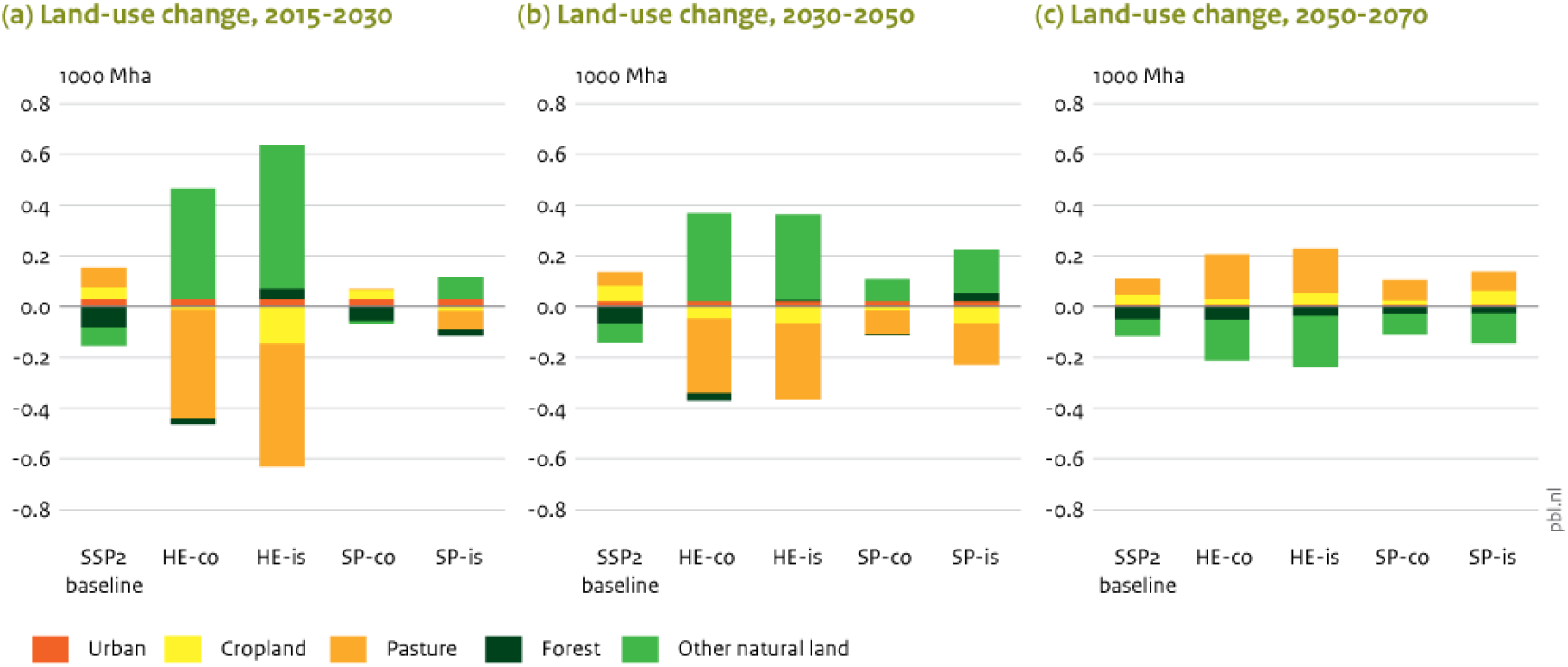
Land use changes in the periods (a) 2015-2030, (b) 2030-2050 and (c) 2050-2070 (SSP-2 baseline, HE-co, HE-is, SP-co, SP-is).

Land-use changes in conservation only scenarios (HE-co and SP-co) contribute only slightly to the reduction of greenhouse gas emissions and to increased carbon sequestration, resulting in limited climate change mitigation. The integrated measures in HE-is and SP-is include large mitigation efforts achieved by industry and energy sectors. Figure 3a shows that the contributions of land-based mitigation in relation to the full mitigation package in HE-is and SP-is will almost double. About 8% of the total results is from implementing bio-energy (van Vuuren et al., 2015), and 4% via technological abatement measures in agriculture such as nitrification inhibitors for N2O from fertilizer application, or farm-scale digesters for CH4 from animal manure (Figure 3a). In the HE-is and SP-is scenarios, emissions from agriculture and forests and other land use sectors are substantially reduced. The cumulative emission reduction from land increases from 4 to 11 and to 13% through integrated measures in HE-is and SP-is scenarios, respectively. In both scenarios, about a quarter of this emission reduction arises from the conservation measures, mostly through avoided deforestation and agroforestry carbon stocks (only SP). These emission reductions could be ‘utilized’ in two ways: to allow a less-dramatic emission decline in energy and industry in the next decades (as illustrated in Figure 3a), or to reach more ambitious climate targets than conventional mitigation projections (RCP 2.6) (Figure 3b). In this latter case, another 0.2 degree below the RCP2.6 scenario (consistent with 2 degrees) might be gained. It must be noted, however, that the land carbon sinks, like afforestation and natural regrowth, are largely temporary, as carbon uptake levels off after some time, depending on the ecosystem.

**FIGURE 3:**
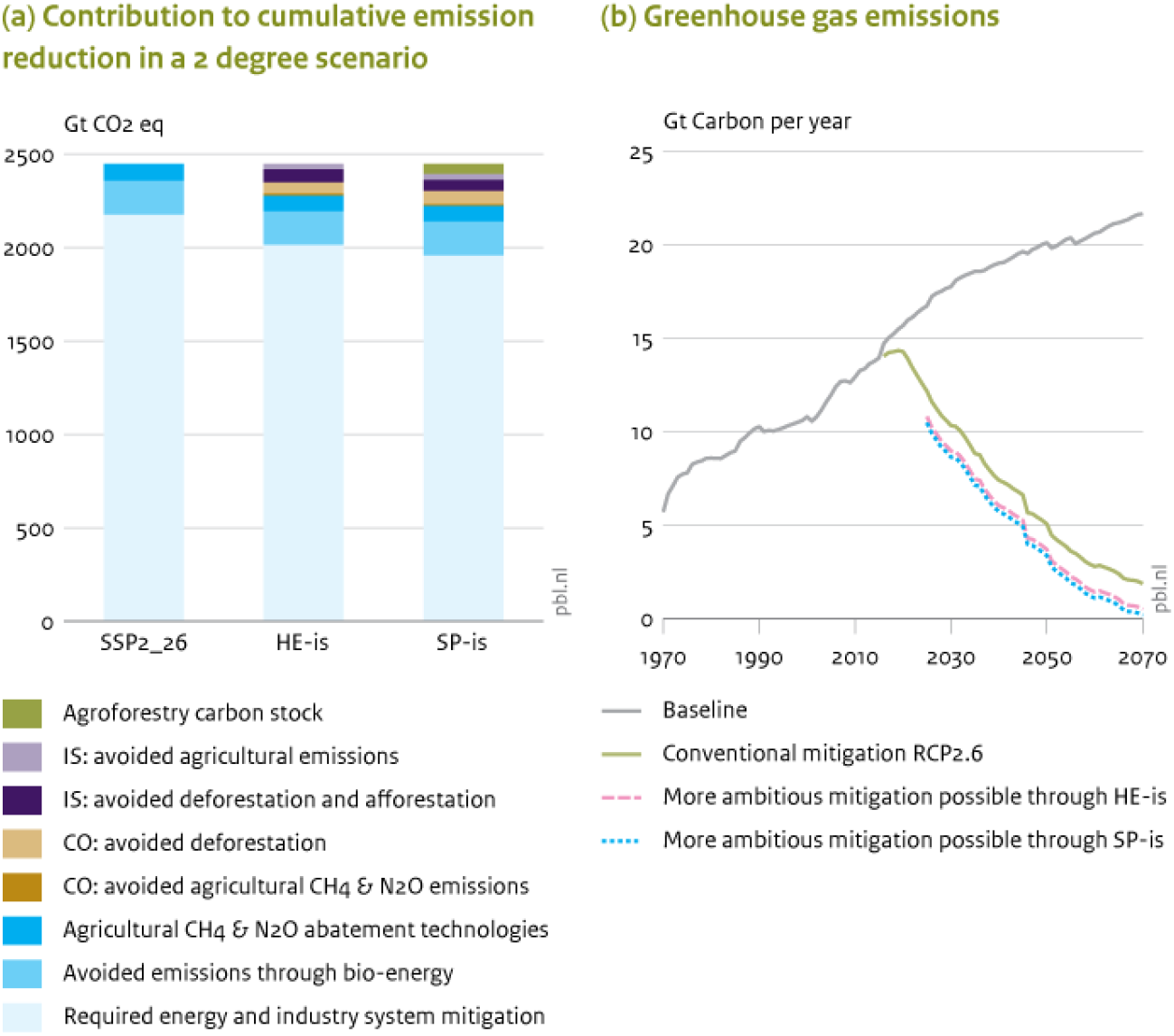
Greenhouse gas (GHG) emission reductions in the different pathways. (a) Contribution of different measures to the required GHG emission reduction in a 2 degree scenario, for conventional mitigation (SSP2 mitigation, based on Van Vuuren et al., 2017), and the HE-is and SP-is scenarios; (b) GHG emissions for baseline and conventional mitigation, and additional mitigation achieved through the HE-is and SP-is scenarios.

In the baseline, between 2015 and 2070, food security increases, as indicated by the fraction of population at risk of hunger (‘prevalence of undernourishment’) being reduced from 10.1 to 2.8% (figure 4a). In conservation only scenarios (HE-co, SP-co) food security risks also reduce, but to a lesser extent, and result in 1.5 to 2 times higher number of people at risk of hunger by 2070 compared to the baseline. The increased scarcity of land, the required intensification (HE), or the shift to agroecological methods (SP) will drive up food prices which reduces access to food for the poor. The integrated sustainability scenarios (HE-is, SP-is) compensate the increased food security risks from the ‘conservation only’ scenarios completely, by reducing both demands for food production and food prices. Food prices are a little lower than to those in the baseline and people at risk of hunger stabilizes just below the baseline SSP2 outcomes, with the SP-is scenario performing better than HE-is. In all scenarios food security risks are most prevalent in Sub-Saharan Africa and South Asia (figure 4b).

**FIGURE 4:**
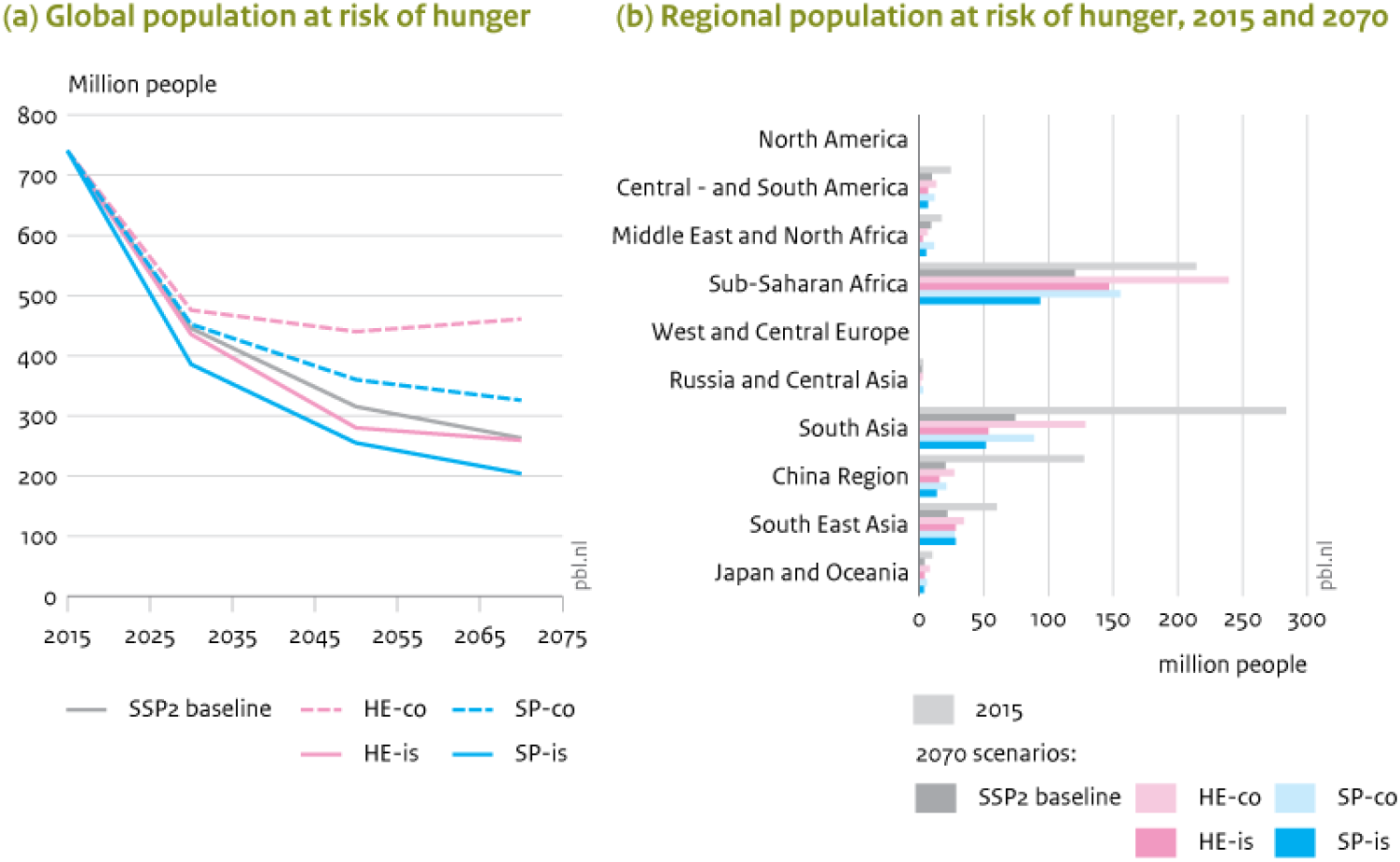
Impacts on food security (a) global and (b) regionally differentiated (Baseline, HE-co, HE-is, SP-co, SP-is).

### 3.3 Projected changes in biodiversity and ecosystem-services

In the baseline scenario, a significant additional loss of biodiversity on land and in freshwater systems is projected (Figures 1 and 5). Both conservation only scenarios (HE-co and SP-co) are able to prevent a large share of the baseline loss of biodiversity. With additional sustainability measures (HE-is and SP-is) global biodiversity shows further improvements as compared to the conservation only scenarios to a level just above 2015 levels in terms of MSA (Figures 1 and 5). Ecosystem services improve especially in the SP scenarios.

**FIGURE 5.**
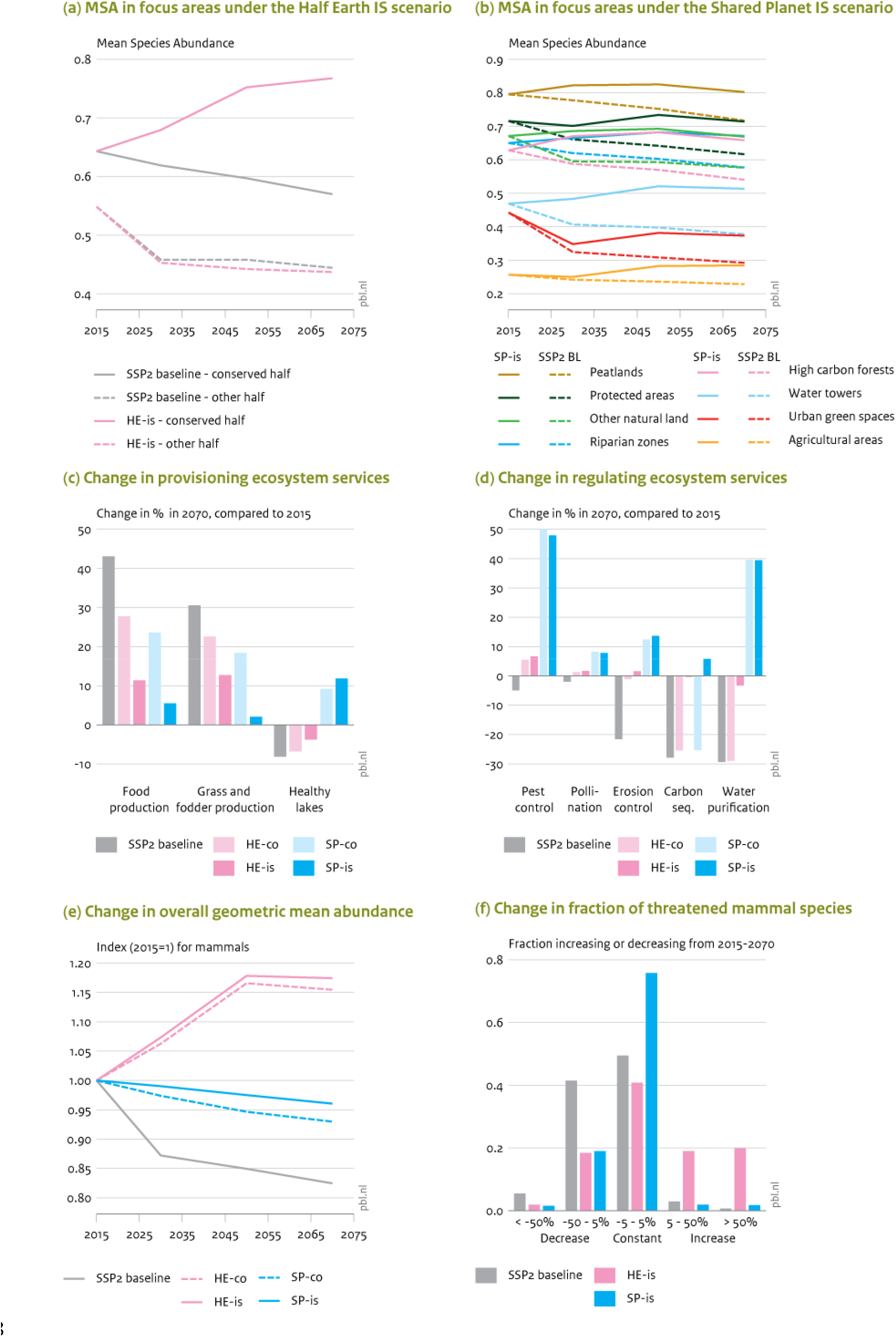
Changes in biodiversity and ecosystem services. (a) MSA under HE-is in protected and in non-protected half; (b) MSA under SP-is for various ecosystems; (5c) Change in provisioning ecosystem services; (c) Change in regulating ecosystem services, 2015-2070; (d) Change in overall geometric mean abundance; (d) Change in fraction of threatened species; 2015-2070.

The biodiversity indicators do, however, respond differently to the two conservation strategies. While both HE-is and SP-is scenarios have similar overall global land MSA results, they show different trends. In HE-is, MSA increases significantly in the protected half, but decreases in the non-protected half, where it follows the baseline value (Figure 5a, see also Figure 6). Conversely, MSA results on land are more equally distributed in the SP-is scenario, indicating that MSA is retained in many regions, and both in natural and human dominated areas. The conservation and restoration of some specific focus areas, supporting ecosystem services, in the SP scenarios will lead to significant increases of MSA in these areas (Figure 5 b, see also Figure 6). The GMA improves in the conservation only scenarios compared to the baseline. This is also reflected by the large number of currently threatened species that increase in population size (Figure 1). The SP strategy, instead, results in a stabilization of the current state, as indicated by the lack of change (or slight decrease) in GMA values compared to current levels and by the fact that most of the threatened species do not change (Figures 5e-f). Note that the impacts of climate change were not included in this analysis, and that the outcomes cover mammals only.

**FIGURE 6.**
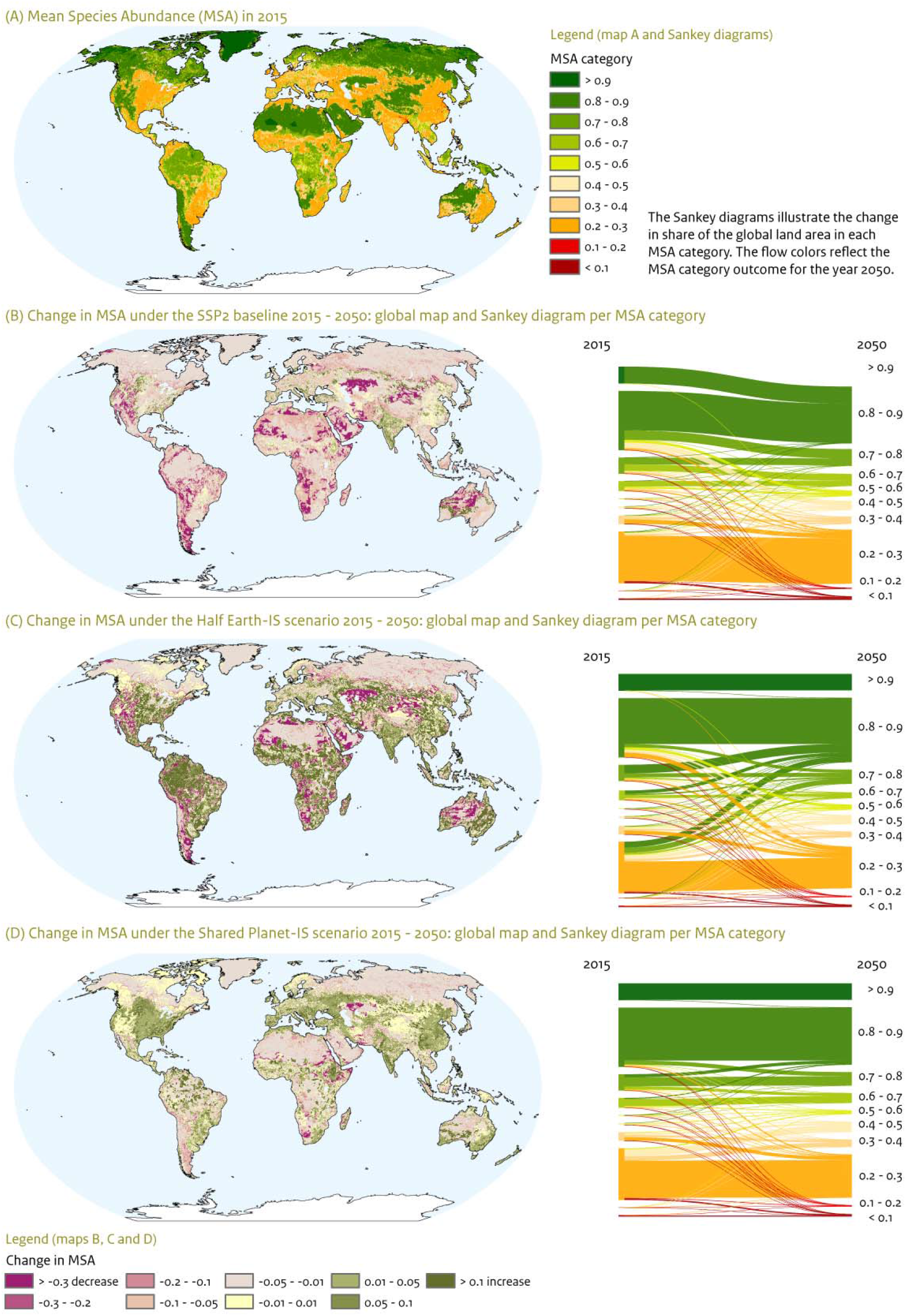
Regional results: (a) Mean Species Abundance in 2015; (b-d) Change in Mean Species Abundance 2015-2050 baseline, HE-is and SP-is. The flow diagrams illustrate the changes in share of global land area for each MSA category between 2015 and 2050.

**FIGURE 7:**
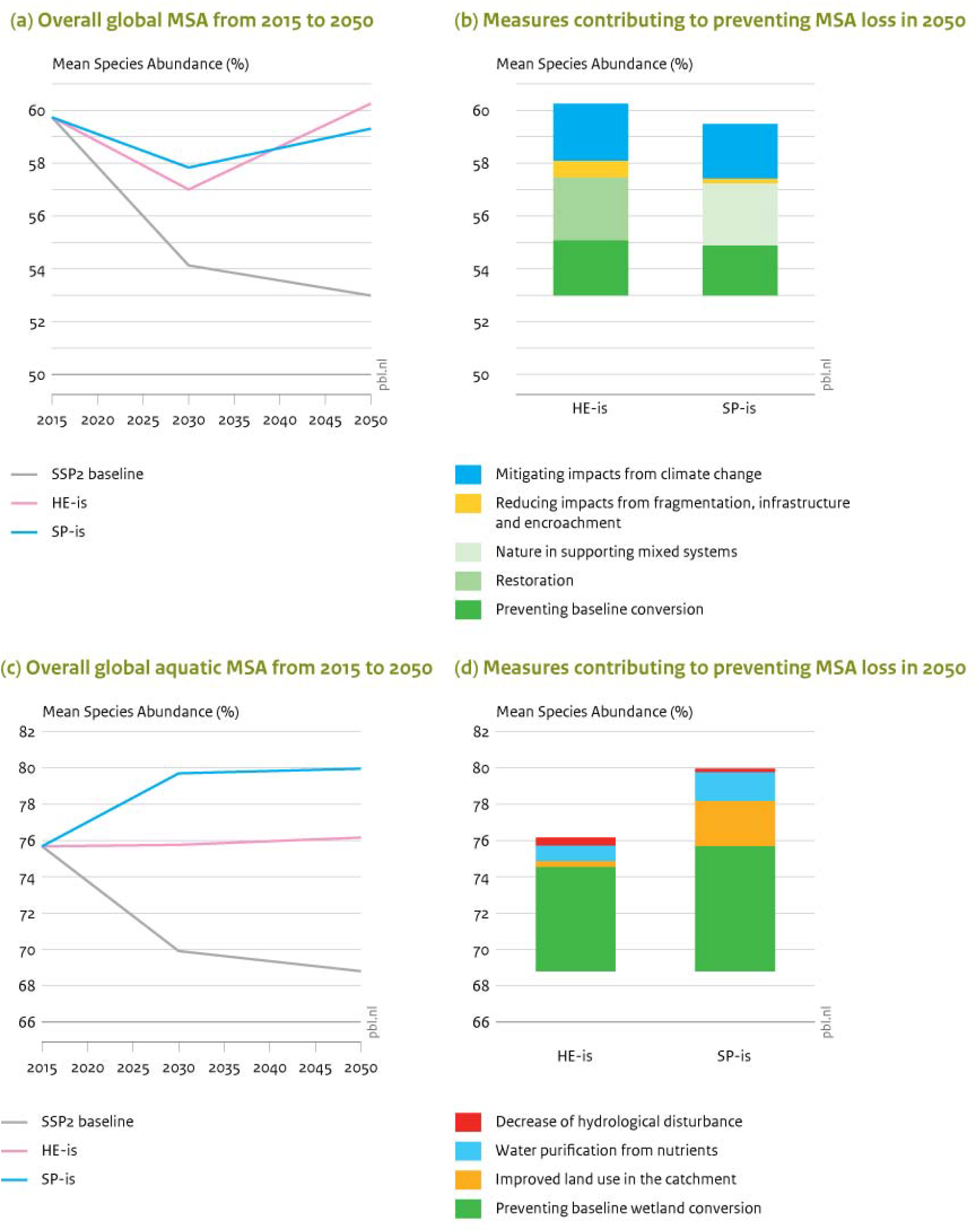
Measures contributing to reducing terrestrial (7.a and b) and aquatic (7.c and d) biodiversity loss, globally (in MSA). Figure 7 describes the net contribution of the various actions in halting the loss and restoring biodiversity compared to the SSP2 baseline outcome over the period 2015 - 2050. The overall difference for the timeline in global MSA between the SSP2 baseline and the HE-is and SP-is is shown for terrestrial and aquatic MSA in figures 7a and 7c. Figure 7b and 7d show how the projected improvement in MSA values in HE-is and SP-is in 2050 compared to the SSP2 baseline is attributed to the various measures in both pathways.

While MSA in water decreases in the baseline, it increases in the SP scenarios and it stabilizes around current values in the HE scenarios (Figure 1). The integrated sustainability scenarios (HE-is, SP-is), result into similar MSA values in water as compared to conservation only scenarios (HE-co, SP-co). This pattern is consistent for the three different water ecosystem types (lakes, rivers and wetlands, but differs between regions (see details in S.5).

In the baseline scenario, provisioning services, such as crop and grass production, increase following agricultural area expansion, but most regulating ecosystem services will decrease, such as erosion control, carbon sequestration and, to a lesser extent, pest control, pollination and the percentage of healthy lakes (figure 1 and 5c-d). This overall pattern in the baseline changes into a pattern of slightly increasing provisioning and clearly increasing regulating services in both HE and SP scenarios, but most pronounced in SP. Most ecosystem services improve due to changing agricultural, forest and water management, however the carbon sequestration relative to the carbon emissions decreases in HE-co and SP-co, and it only shows a slight increase in HE-is and SP-is, as not only the uptake of carbon increases but the emissions decline simultaneously. Overall, most pathways indicate diminishing trade-offs between these two ecosystem services categories (Figure 5c-d). For some variables there are however large regional differences.

The differences between the pathways become more evident when looking at the regional results, as shown in Figure 6 for the changes in MSA on land. MSA in 2015 shows low MSA in human dominated areas and high MSA values in nature dominated parts of the world, including deserts, boreal forest and tundra, and tropical forest areas. The SSP2 baseline results in predominantly decreasing MSA trends in all regions, except for regions where MSA is already and remains low, such as South Asia, Eastern China, Eastern USA and Europe. The largest MSA decreases are expected in regions that currently still show high MSA values and with high demand for land, such as in Sub-Saharan Africa, Central Asia, and the western parts of South and North America, as well as in Middle East and North Africa and western Australia. Figure 6b illustrates the intra-regional dynamics of HE-is scenario. Improvements in MSA resulting from conservation and restoration in the protected areas alternate with large MSA decreases, following the baseline, from conversion of nature in non-protected areas within the same region. Figure 6c shows the SP-is scenario protects important high MSA value areas with pristine nature and results in improving MSA in mixed systems and areas supporting ecosystem services (see also Figure 5b). This, overall limits large MSA decreases and provides a globally more balanced outcome, compared to the HE-is and baseline scenarios. Overall, there are much greater land use shifts projected in the HE and baseline scenarios than in the SP scenario.

Although in HE scenarios all regions are required to achieve high levels of conservation to meet the goals of 50% protection of all ecoregions, efforts differ among regions compared to currently protected areas. Especially in Central and South America a considerable expansion of conserved areas is required (See figure S5.1). In 2050, West and Central Europe will be the region with the lowest percentage change of protected areas compared to current protection level (+17%). The SP scenarios require smaller levels of area conservation than HE scenarios across all regions, with the exception of the South East Asia region that will conserve 51% of its area by 2050. 7 out of 10 regions will expand area conservation by less than +12% compared to the current state. Only the South East Asia (+29%), Central and South America (+19%) and North America (+17%) regions achieve higher levels of area conservation.

### 3.4 Contribution of actions to avoided loss of MSA

Overall, for the global terrestrial MSA in 2050 the projected improvement compared to the baseline is over 7% in the HE-is and over 6% in the SP-is scenario. These improvements can be attributed to several actions resulting from the overall integrated measures on agricultural productivity and agro-ecology, consumption changes and waste reduction. The MSA improvements are the outcome of (1) conservation efforts in order to prevent baseline nature conversion, (2) the large scale rewilding and restoration of natural areas that characterizes HE-is and (3) the improvement of nature in ecosystem services supporting areas and mixed cropland/nature systems that is key in SP-is. Additionally, reducing impacts from (4) fragmentation, infrastructure and encroachment. and climate mitigation measures (5) contribute to the improvement of MSA compared to the baseline outcome in 2050 in both scenarios.

For the global aquatic MSA (area-averaged over the different water types) the projected improvement in 2050 in HE-is and SP-is compared to the baseline, is over 7% and 11% respectively. These improvements can be attributed to: (1) prevented wetland conversion; (2) decreased adverse effects on wetlands by human land-use in their catchment; (3) decreased pollution of lakes and rivers from agricultural and/or urban sources; (4) decreased hydrological disturbance of rivers and floodplain wetlands by restrictions on dams. The effect of climate change in itself is not included because of current model restrictions. The attributions 1-3 contribute more in the SP pathway than in HE, as the former emphasizes the catchment-scale processes and inland wetland conservation for ecosystem services. The contribution of dam restrictions appears to be small on average, but it would be larger if only rivers and floodplain wetlands were considered. This attribution contributes more in HE as more emphasis is given to the conservation of pristine rivers.

## 4 Discussion

### 4.1 Nature, climate and food security

This paper shows that two strong, but contrasting, conservation strategies, Half Earth and Sharing the Planet, combined with an extensive set of sustainability measures, can achieve multiple sustainability goals, including bending the curve for biodiversity, ensuring food security and keeping global temperature change below two degrees. Strong conservation strategies alone will however neither be sufficient to restore biodiversity, as indicated by still decreasing MSA values in both HE-co and SP-co scenarios, nor ensure food security nor mitigate climate change below 2 degrees. Land-use changes in both conservation scenarios (HE-co; SP-co) somewhat contribute to climate change mitigation, but not much, whereas climate change still has impacts on biodiversity.

This confirms the need for strong climate mitigation efforts to be able to bend the curve for biodiversity (Ohashi et al., 2019; Pereira et al., 2010; Soto-Navarro et al., 2020; Warren et al., 2013). So far, current sustainability scenarios such as SSP1 have, however, been insufficient to ‘bend the curve’ for biodiversity loss (Schipper et al., 2020; Pereira et al., 2020; van Vuuren et al., 2015; Kok et al 2018). Leclere et al. (2020, in press) analyzed a very ambitious integrated action portfolio, concluding that strong conservation and food system transformation are key for reversing biodiversity decline. That analysis, however, did not include the impacts of climate change on biodiversity. In this paper, we show that that bending the curve for biodiversity is still possible when taking into account climate change, provided that ambitious climate change mitigation measures are also included. Among these measures, demand reduction measures, will also have added benefits for biodiversity.

To deal with potential trade-offs between biodiversity conservation and climate mitigation, we limited bio-fuels (Behrman et al., 2015; Foley et al., 2011; Raghu et al., 2011) and hydropower use (Ziv et al., 2012; Hermoso, 2017; Reid et al., 2019), did not include climate policy-driven afforestation (Doelman et al., 2019), and maximized the potential gains of land use options (Griscom et al., 2017; Nunez et al., 2020; Roe et al., 2019). Setting limits to applying these energy sources will have consequences for the energy system, such as increased demand for other renewables like solar energy, possibly higher costs, and increased need for improved energy efficiency and reduction of energy demand (van Vuuren et al. 2018).

Furthermore, we show that, both conservation-only strategies and their related agricultural systems will see a slower decrease of number of people at risk of hunger. This confirms possible trade-offs between agricultural production, food security and conservation (Egli et al., 2018; Foley et al., 2011; Mehrabi et al., 2018; Ramankutty et al., 2018) and emphasizes that either of the conservation-only strategies HE-co and SP-co need to be combined with structural changes in both agricultural production and food systems (Brussaard et al., 2010; Fisher et al., 2017; Garnett et al., 2013; Kremen, 2015; Leclere et al. accepted).

Ultimately, to restore nature and achieve food security and climate mitigation targets, both strong area-based conservation and restoration efforts (Dinerstein et al., 2017, 2019; Locke, 2013; Visconti et al., 2019; Wilson, 2016) as well as a broader sustainability measures, including reform of agricultural and food systems, including bothsupply and demand, and other effective climate mitigation, are needed to restore nature (Leclere et al., 2020; Mehrabi et al., 2018; Pereira et al., 2010; Soto-Navarro et al., 2020; Warren et al., 2013). We show that limiting animal consumption, reduction of food waste, and improved agricultural management, to both approaches (sparing and sharing) is key to solve trade-offs between food production and biodiversity conservation, and also a key element of greenhouse-gas emission reduction. This confirms earlier publications on the broad action portfolio required for biodiversity conservation and food security (Foley et al., 2011; Garnett et al., 2013; and Godfray, 2014; Mehrabi et al., 2018; Perfecto & Vandermeer, 2009; van Vuuren et al., 2015, sCBD 2014,). Note, however, that our analysis does not account for socio-economic and political dimensions that have an important role in ensuring food security (Fischer et al., 2017).

### 4.2 Comparing conservation strategies

While the HE and SP strategies show different outcomes with respect to biodiversity, they have also some results in common. Firstly, there is an overlap in conservation areas, inevitably because both strategies include existing protected and key biodiversity areas, but also because they tend to be selected by the ecological criteria in HE (e.g. ecoregion representation, Range Rarity Index) and the ecosystem service criteria in SP (e.g. high-carbon forests, riparian zones). Together the two strategies have over 34 million km2 of conserved area in common, indicating 51% and 80% of the total conserved area for HE and SP respectively. When looking at MSA improvement, in both strategies, 28% of the land area (scattered globally, see Figure SI 5.6 in S.I.) shows at least 5 percent-point improvement.

Secondly, the results for biodiversity and for regulating ecosystem services both point in the same direction, although with differences in magnitude between the strategies. We do not find many trade-offs between biodiversity conservation and regulating ecosystem services. Trade-offs with provisioning services such as crop and grass production do occur (as shown by IPBES, 2019 and Pereira et al., 2020), but they diminish in sustainability scenarios (such as SSP1), and even more in the HE-is and SP-is scenarios we present here. Hence, we show that synergies between the two are certainly possible in important respects, for instance in case of protection of carbon-rich, water-conserving or soil-protecting ecosystems, riparian zones or adopting mixed farming systems. But no doubt, in specific cases trade-offs between conservation goals and ecosystem services will occur (Maes et al., 2012; Van der Biest et al., 2020).

There are, however, also large differences between the HE and SP strategies. HE includes ~20% extra conservation areas compared to SP, while SP includes a number of biodiversity improvements in agricultural areas. On average, HE performs better than SP on most terrestrial biodiversity measures, but this varies among indicators and among regions. In general, HE greatly improves biodiversity in the protected half but has hardly any effect in the non-protected half, while SP shows more evenly distributed improvements. Especially the Amazon and Cerrado regions, Central Africa and South and Southeast Asia improve most in HE-is. Some other regions score better MSA (hence biodiversity) results under the SP-is scenario, such as the western part of North America, central and east Asia, western Australia and scattered parts of Africa and South America. It appears that HE (compared to SP) conservation strategies do not necessarily translate into higher regional MSA results, thereby suggesting that a protection based primarily on wilderness might not be the optimal solution everywhere (Ellis & Mehrabi, 2019; Pimm et al., 2018). Nevertheless, our results are consistent with the conclusion by Riggio et al. (2020) that there are still low-impact areas available in the world that provide opportunities for conservation. This is also in line with recent criticisms by Visconti et al. (2019) and Jung et al. (2020) who, like Pimm et al., (2018), point at the importance of allocating the right places for conservation. Our analysis suggests that a more differentiated approach is needed to strike optimal results across different regions and the projected MSA improvement in a region will depend on the initial MSA value (i.c. in 2015) in that region. Our findings align well with the concept of the ‘Three Conditions’ as presented by Locke et al. (2019) that takes a middle position between HE and SP. The ‘Three Conditions’ identify different regional conditions ranging from large wild areas, via shared lands to cities and farmland, to enable the development of conservation responses and production practices appropriate for each condition that can be informed by both HE and SP.

Another difference between the two conservation pathways is that aquatic MSA in general profits more from the SP strategy than terrestrial MSA. This may be attributed to the fact that many aquatic ecosystems are more connected with each other and with the surrounding or upstream land, e.g. by flow of water and substances. This implies that general improvements in land management, including the restoration of riparian buffer zones, have a broader effect than only locally, while on the other hand local conservation measures might be less effective for aquatic ecosystems than for terrestrial ones, unless whole landscapes or (sub)catchments are protected. This is in accordance with the emphasis on catchment-based conservation measures for aquatic biodiversity recently put forward by Tickner et al (2019) and Van Rees et al (2020). In addition, added benefits would be expected in the SP strategy from the lower use of agrochemicals. On the contrary, HE scores better on the effect of flow on biodiversity as more emphasis is put on river flow restoration (another recommendation by Tickner et al (2019) and Van Rees et al (2020) due to more restrictions on hydropower. HE would probably also score better for water bodies with many endemics (‘reversed islands’), but this could not be explored in this study as specific species models were not applied.

Comparing both conservations strategies with respect to land cover types, the HE strategy, requires substantial changes in land use, especially grazing lands are expected to be abandoned and restored towards natural grasslands, such as savannah and steppe systems, while there is less dynamics in forest systems (see Figure 2). With respect to species, HE protects more areas important for the conservation of threatened species, which yields decreasing number of species at risk of extinction and higher RLI values, whereas in SP the human dominated landscapes will alter considerably, improving the status of species that are adapted to these landscapes. This shows the value of using different biodiversity indicators in parallel, the GMA and RLI providing information on species trends and the MSA more on the ecosystem level.

A consequence of the large expansion of conserved areas, the HE scenario will directly affect a large amount of people, and this could put nature against humans risk diminished acceptance of nature conservation (Buscher & Fletcher, 2020; Ellis, 2019; Schleicher et al., 2019). Furthermore, a too-narrow focus on achieving quantitative protection targets might not necessarily ensure that biodiversity is actually conserved (Visconti et al., 2019). Attention should also be focused on the management and governance of the conserved areas so as to ensure fair and effective protection.

### 4.3 Actions and efforts needed

The scenarios show that a portfolio of measures is needed to achieve nature, food and climate targets in both the HE and SP conservation strategies. A transition towards more sustainable forms of agriculture is a cornerstone for all strategies to halt biodiversity loss (Gross et al., 2019; Kremen, 2015; Phalan et al., 2011). Earlier analyses showed the importance of changes in agricultural production to create space for area-based conservation and restoration (Kok et al., 2018; Leclere et al., 2018, 2020; Mehrabi et al., 2018). Whether this is implemented by additional technological intensification (Egli et al., 2018; Phalan et al., 2011) as in HE-is, or by a shift to agro-ecological methods (Kremen, 2015, Tittonel, 2014) as in SP-is, both imply considerable changes in agricultural systems. Optimal strategies for agriculture will differ regionally and can be aligned with different kinds of conservation priorities (Locke et al., 2019).

This points at the need for improved spatial planning approaches to allocate conservation areas; not only in the traditional form of Protected Areas, but also in new forms of conservation, such as the recently internationally agreed Other effective Conservation Measures (Dudley et al., 2018; Garnett et al., 2018; IUCN WCPA, 2019). Also new proposals for conservation are being suggested such as ‘Promoted Areas’ (Buscher & Fletcher, 2020). Improved spatial planning and management of conservation areas may enhance the effectiveness of conservation areas also for freshwater biodiversity (Hermoso et al., 2016; Reis et al., 2017; van Rees et al., 2020). All emphasize sustainable use and the delivery of socially, economically and culturally relevant benefits next to conservation objectives. The SP strategy comes close to these ideas where the implementation of a landscape approach delivers conservation and Nature’s Contribution to People (NCP) at the same time (Kremen & Merenlender, 2018).

Crucially important elements in a portfolio of measures are reducing food waste (Kok et al., 2018; Parfitt et al., 2010; Tscharntke et al., 2012) and reduced overall consumption of animal products, (Machovina et al., 2015; Stoll-Kleemann & Schmidt, 2016; Tilman & Clark, 2014, Springmann, 2018; Willet et al., 2019, FAO et al, 2020). Improved human health and more attention to animal welfare are important co-benefits of such diets that can be used as leverages for implementation. Reducing food waste is also fundamental as according to FAO (2013) one third of produced food for human consumption is lost. As food waste happens at different point of the supply chain for different regions (low-income at agricultural production; middle-high income at retailer and consumer level) (FAO, 2013; Tscharntke et al., 2012) this requires multiple interventions.

Effective climate change mitigation is an important condition to restore nature, especially in the long term (after 2050) (e.g. Reid et al., 2019.) We have also shown that conservation measures can contribute considerably to up to 134 and 309 Gt CO-2 equivalents (6 and 13% %) of required GHG reductions, for conservation only and integrated strategies, respectively. This is lower than recent estimates on the contribution of land or nature-based climate solutions to climate (Griscom et al., 2018; Nunez et al., 2020; Roe et al., 2019). This can be explained by the fact that our study focusses on biodiversity-related measures, and does not cover all land-related measures in agriculture, grassland and wetlands, and afforestation as included by Griscom et al. (2018). The recent discussion on the potential of afforestation (Bastin et al., 2019) triggered worries about delayed action in the energy sector (Friendlingstein et al., 2019), and strong land-based mitigation may also affect food systems. (Doelman et al., 2018). Furthermore, carbon uptake through restoration measures is effective predominately in the short-term, though some long-term sinks exist in natural systems.

### 4.4 Methodological reflections

The analysis in this paper is based on the IMAGE and GLOBIO models. This allowed us to analyse multiple pressures on biodiversity, as well as multiple indicators for biodiversity and ecosystem services in both the terrestrial and freshwater domain. The analysis covers the most important drivers and pressures on the global scale, but is not complete, as for instance the influence of invasive species and of toxic stress are not included. Likewise, feedbacks of ecosystem services to production sectors are not implemented, and the modelling of mixed farming systems could be further improved. It would also be important in a next step to analyse these pathways using multiple models. Although we did not explicitly analyse model- and scenario-uncertainties for this study, together the two strategies span a solution space for possible pathways reflecting major uncertainties. Projecting the scenarios towards 2070, thus going beyond the 2050 target year of the CBD vision, shows that in the longer term integrated sustainability measures taken in the scenarios are not enough to compensate for the projected growth in population and wealth in the baseline. This again emphasizes the importance to address the indirect drivers of biodiversity loss such as economy and population, as well as institutions, equity aspects and underlying value systems (IPBES, 2019; Buscher et al., 2017; Buscher & Fletcher, 2020; KC and Lutz, 2017; Otero et al., 2020; Pascual et al., 2017; Perfecto & Vandermeer, 2009; Tscharntke et al., 2012).

So far, explorative scenario-analysis have mainly looked at achieving biodiversity objectives from a climate perspective (Pereira et al., 2020; Riahi et al., 2017), and show the continued decline of biodiversity. In this study, we bring ambitious biodiversity, together with climate objectives in scenario-analysis (Dinerstein et al., 2019; Pereira et al., 2017, Titeux et al., 2017, Warren et al., 2013; Soto-Navarro et al., 2020), and to put nature and our relationships with it at the center of biodiversity scenarios (Rosa et al., 2017). As shown, this requires inclusion of important considerations in scenarios such as criteria for conservation areas (Riggio et al., 2020), ecosystem services (Soto-Navarro et al., 2019) or nature-inclusive forms of agriculture (Bommarco et al., 2018).

We aimed to build upon and contribute to the work of IPBES and their recent Nature Futures Framework (IPBES-NFF) which provides guidance to elaborate scenarios with different types of human-nature relations at the centre (Pascual et al., 2017; Rosa et al., 2017; Pereira et al., 2020). Specifically, the HE scenario was envisioned to prioritize the intrinsic value of nature as it aims at conserving nature’s diversity and functions (‘Nature for Nature’ according to IPBES-NFF). The SP scenario was envisioned to prioritize instrumental and relational values, (‘Nature for Society’ and ‘Nature as Culture’). However, due to limitations in our models, only the ‘Nature for Society’ part could be operationalized into the model. Efforts are underway that aim to include Nature as Culture in global scenario-analysis. The way forward may be through a combination of quantitative and more qualitative methods. The multiple ways people relate to and value nature could, in this way, become evident and recognized in a conservation context (Chan et al., 2016; Martin et al., 2016; Pascual et al., 2017, van Zeijts et al., 2017). Better inclusion of a plurality of values in scenario-analysis would also help to better understand the equity dimensions of conservation, for example for people living in or around the conservation areas, as put forward by Schleicher et al. (2019) and Büscher et al. (2017), and to reconcile the conservation goals with local or regional benefits.

### 4.5 Implications for post-2020 Global Biodiversity Framework

The analysis in this paper has various implications for the post-2020 Global Biodiversity Framework that is currently under negotiation. First and foremost, to meet long term goals for biodiversity, climate change, food security (CBD, UNFCCC, SDGs), it will be necessary to advance a portfolio of ‘integrated sustainability measures’, that combines strong conservation efforts with addressing direct and indirect drivers of biodiversity loss, especially land use change, agriculture and climate change and increased attention to capturing the potential of nature’s potential contribution to resolving global challenges. These measures include strong area-based conservation and restoration (Dinerstein et al., 2017, 2019; Locke, 2013; Visconti et al., 2019; Wilson, 2016), changes in agricultural production and food consumption (e.g., reducing food waste, reducing animal product consumption and increasing agricultural productivity) (Machovina et al., 2015; Stoll-Kleemann & Schmidt, 2016; Tscharntke et al., 2012), as well as decarbonizing the energy system in ways that are not detrimental to biodiversity and food security and benefit as much as possible from natural climate solutions.

Conservation only efforts, without changes in consumption and waste and additional effective climate mitigation, although resulting in avoiding a considerable part of the projected loss in biodiversity, will not be able to restore nature and have a strong trade-off with food security (Mehrabi et al., 2018). Furthermore it will be important to capitalize on the possible synergies between conservation and climate mitigation policy by indicating the contribution strong conservation efforts can make to bringing climate targets within closer reach (Dinerstein et al., 2019), as well other contributions of nature to people (e.g. agriculture, clean water) (Diaz et al., 2018).

The two conservation strategies not only have different objectives but, in fact, show the effects of protecting different types of nature over other and can be related to the goals and targets currently under negotiation for the post-2020 Global Biodiversity Framework (CBD, 2020). The proposed target of 30% protection by 2030 is coherent with both of the long term strategies analyzed, although each may differ in the prioritization of areas to conserve. Evidently, the two strategies will lead to two alternative futures with different urban, agricultural, forestry and fisheries systems. We do not to advocate one strategy or the other, or present these as irreconcilable pathways, but rather we present these findings to help open up a space where insights from different alternatives can be deliberated.

## Author contributions

MK, RA, JM, JJ designed the research; MK, JM, MI, RA developed scenario-narratives and conservation maps; JM, JH, JJ, MB, AS, RA ran GLOBIO; WJ, ES, AT ran IMAGE; all authors contributed to analysis of results; MK, RA wrote first versions of this paper to which all authors contributed in various iterations, MK and JM finalized the paper.

Authors declare no conflict of interest.

